# Microbial Species Abundance Distributions Guide Human Population Size Estimation from Sewage Microbiomes

**DOI:** 10.1101/2020.12.15.390716

**Authors:** Lin Zhang, Likai Chen, Xiaoqian Yu, Claire Duvallet, Siavash Isazadeh, Chengzhen Dai, Shinkyu Park, Katya Frois-Moniz, Fabio Duarte, Carlo Ratti, Eric J. Alm, Fangqiong Ling

## Abstract

The metagenome embedded in urban sewage is an attractive new data source to understand urban ecology and assess human health status at scales beyond a single host. Analyzing the viral fraction of wastewater in the ongoing COVID-19 pandemic has shown the potential of wastewater as aggregated samples for early detection, prevalence monitoring, and variant identification of human diseases in large populations. However, using census-based population size instead of real-time population estimates can mislead the interpretation of data acquired from sewage, hindering assessment of representativeness, inference of prevalence, or comparisons of taxa across sites. Here, we show that taxon abundance and sub-species diversisty in gut-associated microbiomes are new feature space to utilize for human population estimation. Using a population-scale human gut microbiome sample of over 1,100 people, we found that taxon-abundance distributions of gut-associated multi-person microbiomes exhibited generalizable relationships with respect to human population size. Here and throughout this paper, the human population size is essentially the sample size from the wastewater sample. We present a new algorithm, MicrobiomeCensus, for estimating human population size from sewage samples. MicrobiomeCensus harnesses the inter-individual variability in human gut microbiomes and performs maximum likelihood estimation based on simultaneous deviation of multiple taxa’s relative abundances from their population means. MicrobiomeCensus outperformed generic algorithms in data-driven simulation benchmarks and detected population size differences in field data. New theorems are provided to justify our approach. This research provides a mathematical framework for inferring population sizes in real time from sewage samples, paving the way for more accurate ecological and public health studies utilizing the sewage metagenome.

**Author summary:** Wastewater-based epidemiology (WBE) is an emerging field that employs sewage as aggregated samples of human populations. This approach is particularly promising for tracking diseases that can spread asymptomatically in large populations, such as the COVID-19. As a new type of biological data, sewage has its own unique challenges to utilize. While wastewater samples are usually assumed to represent large populations, it is not guaranteed, because of stochasticity in toilet flushes; unlike epidemiological experiments collecting data from individuals, sample size, i.e., the human population size represented by a wastewater sample, is a fundamental yet difficult-to-characterize parameter for sewage samples. Researchers would need to aggregate data from large areas and week-long collection to stabilize data, during which, important spikes in small areas or short time scales may be lost. It also remains challenging to turn viral titers into case prevalences, evaluating representativeness, or comparing measurements across sites/studies.

This study provides a framework to estimate human population size from sewage utilizing human gut-associated microorganisms. Through analysis, we demonstrate that variance of taxon abundances and single-nucleotide polymorphism as two variables that change with population size. We provide a new tool MicrobiomeCensus that performs population size estimation from microbial taxon abundances. MicrobiomeCensus outperforms generic algorithms in terms of computational efficiency while at comparable or better accuracy. Using MicrobiomeCensus, we detected population size differences in sewage samples taken in Cambridge, MA, under two sampling approaches, i.e., “grab” or “composite” sampling. This study provides a framework to utilize individual-level microbiomes to learn from sewage, paving the way to prevalence estimation and improved spatio-temporal resolutions in WBE..

## Introduction

The metagenome embedded in urban sewage is an attractive new data source to understand urban ecology and assess human health status at scales beyond a single host[22, 2, 29]. Sewage microbiomes are found to share a variety of taxa with human gut microbiomes, where the baseline communities are characterized by a dominance of human-associated commensal organisms from the *Bacteroidetes* and *Firmicutes* phyla[29, 22, 24]. Human viruses like SARS-CoV-2 and polioviruses were detected in sewage samples during the pandemic and silent spreads, respectively, and found to correlate to reported cases, suggesting that sewage samples could be useful for understanding the dynamics in the human-associated symbionts at a population level[25, 21]. Sewage has several advantages as samples of the population’s collective symbionts. For instance, sewage samples are naturally aggregated, wastewater infrastructures are highly accessible, and data on human symbionts can be collected without visits to clinics, thus utilizing sewage samples can reduce costs and avoid biases associated with stigma and accessibility[2, 28]. Consequently, SARS-CoV-2 surveillance utilizing sewage samples are underway globally and incorporated into the U.S. Centers for Disease Control and Prevention surveillance framework[9].

A pressing challenge in utilizing sewage for ecological and public health studies is the lack of methods to directly estimate human population size from sewage. Specifically, virus monitoring at finer spatial granularity, e.g., single university dorms and nursing homes, are informative for guiding contact tracing and protecting populations at higher risk, but real-time population size estimations at such fine granularity are not yet available. For a given area, the census population (*de jure* population) can be larger than the number of people who contributed feces to sewage at a given time (*de facto* population)[11]. Conversely, the *de jure* population can also be smaller than the *de facto* population due to the presence of undocumented individuals[13]. Population proxies that are currently used for monitoring at wastewater-treatment plants, such as the loading of pepper mild mottle viruses, likely have high error at the neighborhood level because of their large variability in human fecal viromes (10^6^-10^9^ virions per gram of dry weight fecal matter) [38]. Consequently, it is difficult to assess the representativeness of a sewage sample, infer the taxon abundance differences across time and space, or interpret errors. Lack of population size information could lead to false negatives in assessing virus eradication, because an absence of biomarkers might be caused by a sewage sample that under-represents the population size. Despite its importance, few studies have explicitly explored ways to estimate real-time human population size from sewage samples independent from census estimates[37].

Macroecological theories of biodiversity may offer clues to decipher and even enumerate the sources of a sewage microbiome. While we are only beginning to view sewage as samples of human symbionts beyond one person, generating multi-host microbiomes resembles a fundamental random additive process. Sizling et al. showed that lognormal SADs can be generated solely from summing the abundances from multiple non-overlapping sub-assemblages to form new assemblages[35]. Likewise, adding multiple sub-assemblages can also give rise to common Species-Area Relationships[35]. For microbial ecosystems, Shoemaker et al. examined the abilities of widely known and successful models of SADs in predicting microbial SADs and found that Poisson Lognormal distributions outperformed other distributions across environmental, engineered, and host-associated microbial communities, highlighting the underpinning role of lognormal processes in shaping microbial diversity[34].

In this study, we conceptualize a sewage microbiome as a multi-person microbiome, where the number of human contributors can vary. We hypothesize that the species abundance distribution in the multi-person microbiome will vary as a function of the human population size, which would arise from summing taxon abundances from multiple hosts analogous to the Central Limit Theorem. We use human gut microbiome data comprising over a thousand human subjects and machine learning algorithms to explore these relationships. Upon discovering a generalizable relationship, we develop MicrobiomeCensus, a nonparametric model that utilizes relative taxon abundances in the microbiome to predict the number of people contributing to a sewage sample. MicrobiomeCensus utilizes a multivariate *T* statistic to capture the simultaneous deviation of multiple taxa’s abundances from their means in a human population and performs maximum likelihood estimation. We provide proof on the validity of our approach. Next, we examine model performance through a simulation benchmark using human microbiome data. Last, we apply our model to data derived from real-world sewage. Our nonparametric method does not assume any underlying distributions of microbial abundances and can make inferences with just the computational power of a laptop computer.

## Results

### Species abundance distributions of multi-person microbiomes vary by population size

We consider the fraction of microorganisms observed in sewage that are human-associated anaerobes as an “average gut microbiome” sampled from residents of a catchment area. Hence, our task becomes to find the underlying relationship between the number of contributors and the observed microbiome profiles in sewage samples. We define an “ideal sewage mixture” scenario to illustrate our case, where the sewage sample consists only of gut-associated microorganisms and is an even mix of n different individuals’ feces (Figure 1). We denote the gut microbiome profile of an individual as *X_i_* = (*X*_*i*,1_ *X*_*i*,2_, …, *X_i,p_*)^⊤^, where each *X_i,j_* represents the relative abundance of operational taxonomic unit (OTU) *j* from individual *i*. Hence our ideal sewage mixture can be represented as

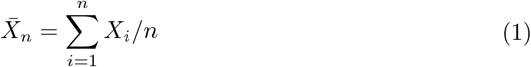

where vectors 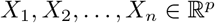 are microbiome profiles from individuals 1, …, *n*. Under the ideal sewage mixture scenario, if we can quantitatively capture the departure of the sewage microbiome profile from the population mean of the human gut microbiomes of people constituting the catchment area, we will be able to estimate the population size.

**Figure 1.**
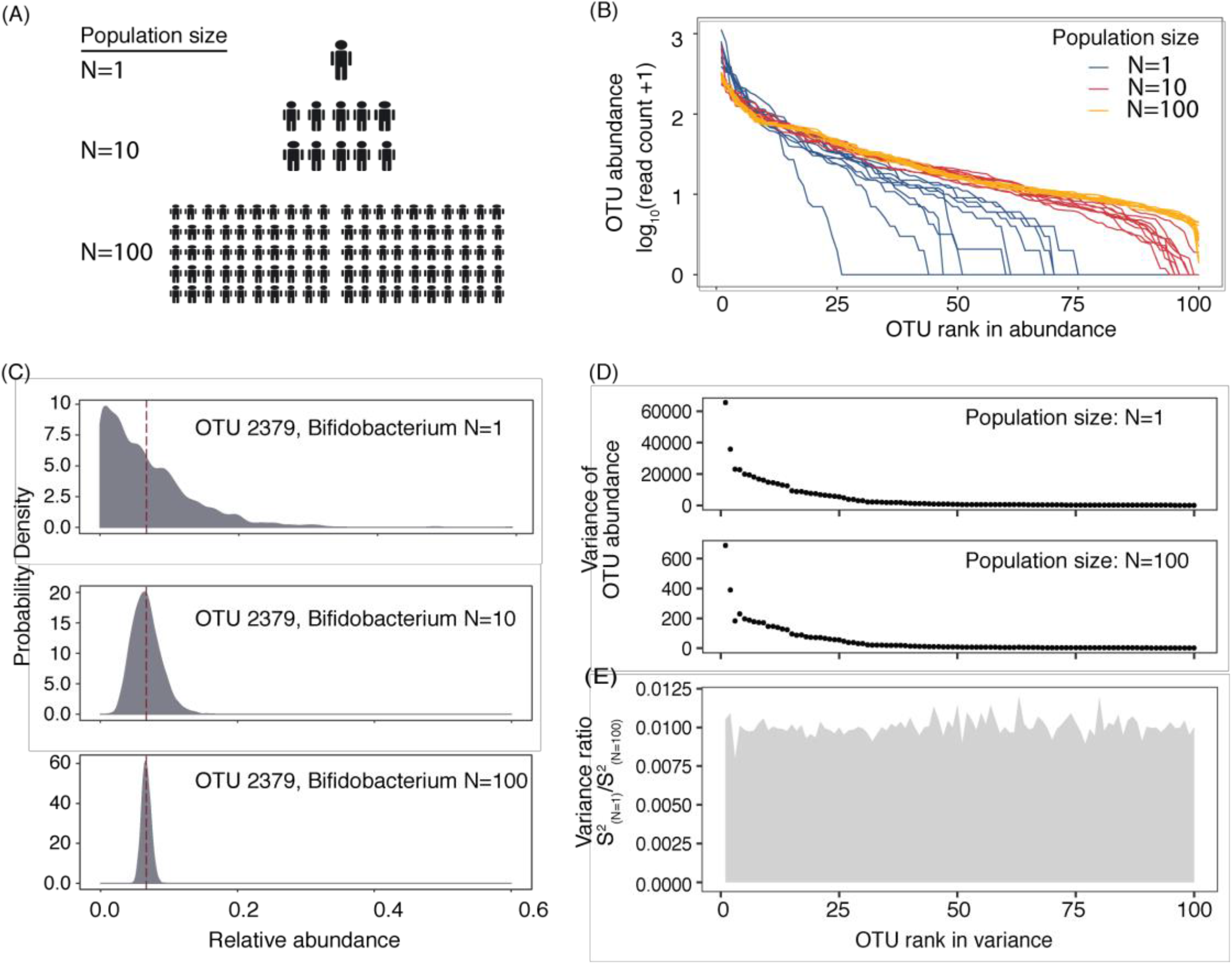
An ideal sewage mixture simulation shows the potential of microbiome taxon abundance profiles as population census information sources. (A) We generated an “ideal sewage mixture” consisting of gut microbiomes from different numbers of people. (B) Ranked abundance curves for gut microbiomes of one person and mixtures of multiple people exhibit different levels of dominance and diversity. Blue lines show the rank abundance curves in stool samples (one person), red lines show 10-person mixtures, and saffron lines show 100-person mixtures. In each scenario, ten examples are shown. All samples were rarefied to the same sequencing depths (4,000 seqs/sample). (C) The probability density function of the relative abundance of one taxon for different population sizes. OTU-2379, a *Bifidobacterium* taxon, was used as an example. Maroon dashed lines indicate the sample means. (D) Multiple taxa’s abundance variances in one-person samples and 100-person samples. The dominant taxa are shown (top100) and are sorted by their ranks in variance. (E) The ratios of the variances of one-person samples and 100-person samples across dominant gut microbial taxa.

Using a dataset comprised of 1,100 individuals’ gut microbiome taxonomic profiles[39], we created synthetic mixture samples of different numbers of contributors through bootstrapping (Figure 1A). First, examined from an ecological perspective, the shape of the ranked abundance curves of the gut microbiomes differed when the means of multiple individuals were examined: when the number of contributors increased, a normal distribution appeared(Figure 1B). For the single-person microbiomes, log-series and lognormal distributions explained 94% and 93% of the variations in the SADs, respectively, compared with 89% for Poisson lognormal, 87% for Zipf multinomial and 80% for the broken-stick model. Multi-person microbiomes were best predicted by log-series or lognormal models, but as the population increased to over a hundred, the multi-person SADs were best described by only lognormal SADs (Table S1).

We explored the distributions of the relative abundances of gut bacteria as a function of population size. As expected, the distribution of a taxon’s relative abundance changes with population size (Figure 1C). For instance, for OTU-2397, a *Bifidobacterium* taxon, the relative abundance distribution was approximately log-normal when the relative abundance in single-host samples was considered, yet converged to a Normal distribution when mixtures of multiple hosts were considered. Although the means of the distributions of the same taxon under different population sizes were close, the variation in the data changed. A smaller variance was observed when the number of contributors increased (Figure 1D). Notably, different taxa varied in the rates at which their variances decreased with population size (Figure 1E), suggesting that a model that considers multiple features would be useful in predicting the number of contributors.

### Classifiers utilizing microbial taxon abundance features alone detects single-person and multi-person microbiomes

Inspired by the distinct shapes of SADs in multi-person gut-microbiomes from those of single-person microbiomes, we set up a classification task using the taxon relative abundances to separate synthetic communities constituting one, ten, and a hundred people. With algorithms of varying complexity, namely Logistic Regression (LR), Support Vector Machine (SVM), and Random Forest (RF) classifier, classification accuracies of 29.6%, 97.2%, and 100% were achieved (Figure 2). Between RF and SVM, RF showed higher sensitivity and specificity in classifying all population groups (Table S2). This experiment suggests the usefulness of microbiome features in predicting human population counts from mixture samples.

**Figure 2.**
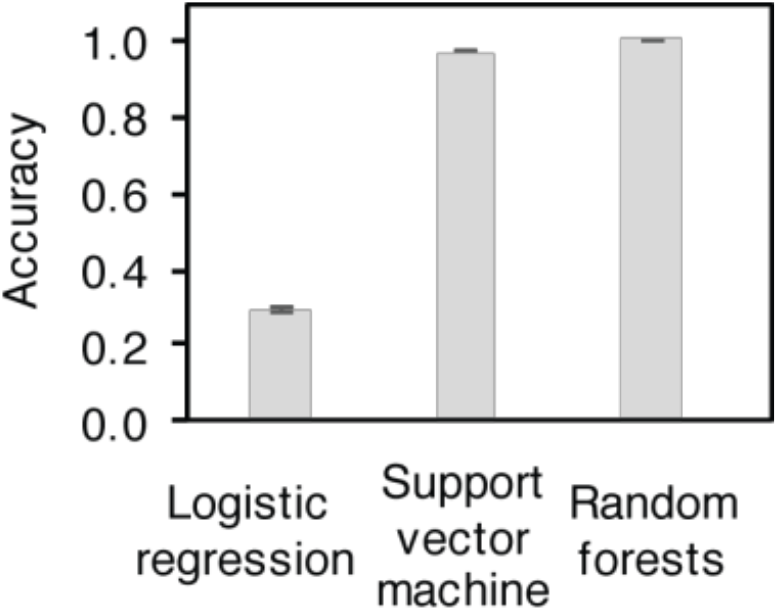
Classifier performance of models utilizing gut microbiome taxon abundances.

### MicrobiomeCensus is a statistical model that estimates population size from microbial taxon abundances

While the classification tasks described above demonstrated the usefulness of taxa’s relative abundances in predicting the population size, a complex model like RF provided little explanatory power. We then ask, since the variance in the relative abundance of a given taxon decreases with population size, can we devise a statistic that captures the simultaneous deviation of several taxa’s abundances from their means, and estimate population size utilizing the statistic? Further, will this new method perform well despite inter-personal variation in gut microbiomes?

Our new method, MicrobiomeCensus, involves a statistic *T_n_* to capture the simultaneous deviation of multiple taxa’s abundances from their means in relation to the variance of those taxa in the population (Figure 3A). We denote 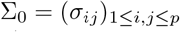 as the covariance matrix for the individual microbiome profile and let Λ_0_ be a diagonal matrix with 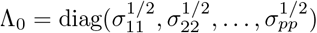. Then the statistic takes form

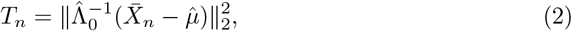

where 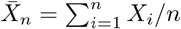 denotes the observed microbiome profile in ideal sewage, *μ* represents the population mean for the catchment area and 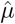 is an estimator, 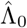 is an estimator of Λ_0_ and 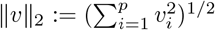 for any vector 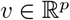. This statistic is enlightened by the classical Hotelling *T*^2^ statistic[16] 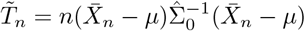, where 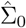 is the sample covariance matrix, an estimator of ∑_0_. Actually if we assume the covariance matrix is diagonal (no correlations between different taxa), then they are essentially the same statistic in view of 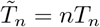. The reason we replace covariance matrix ∑_0_ by its diagonal Λ_0_ is because for high dimensional situations, it would be very difficult to estimate the covariance matrix. In cases when *p* > *n*, the sample covariance matrix is singular and thus 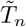 is not even well defined. Studies accommodating the Hotelling *T*^2^ type statistic into the high-dimensional situation can be found, for example, in Bai and Saranadasa[1], Chen and Qi[10], Xu et al[36], etc. Our proposed statistic can handle the high dimensional cases as well, since the diagonal entities Λ_0_ can be well estimated even when *p* is large. And we extend its application beyond the problem of the significance of the multivariate means.

**Figure 3.**
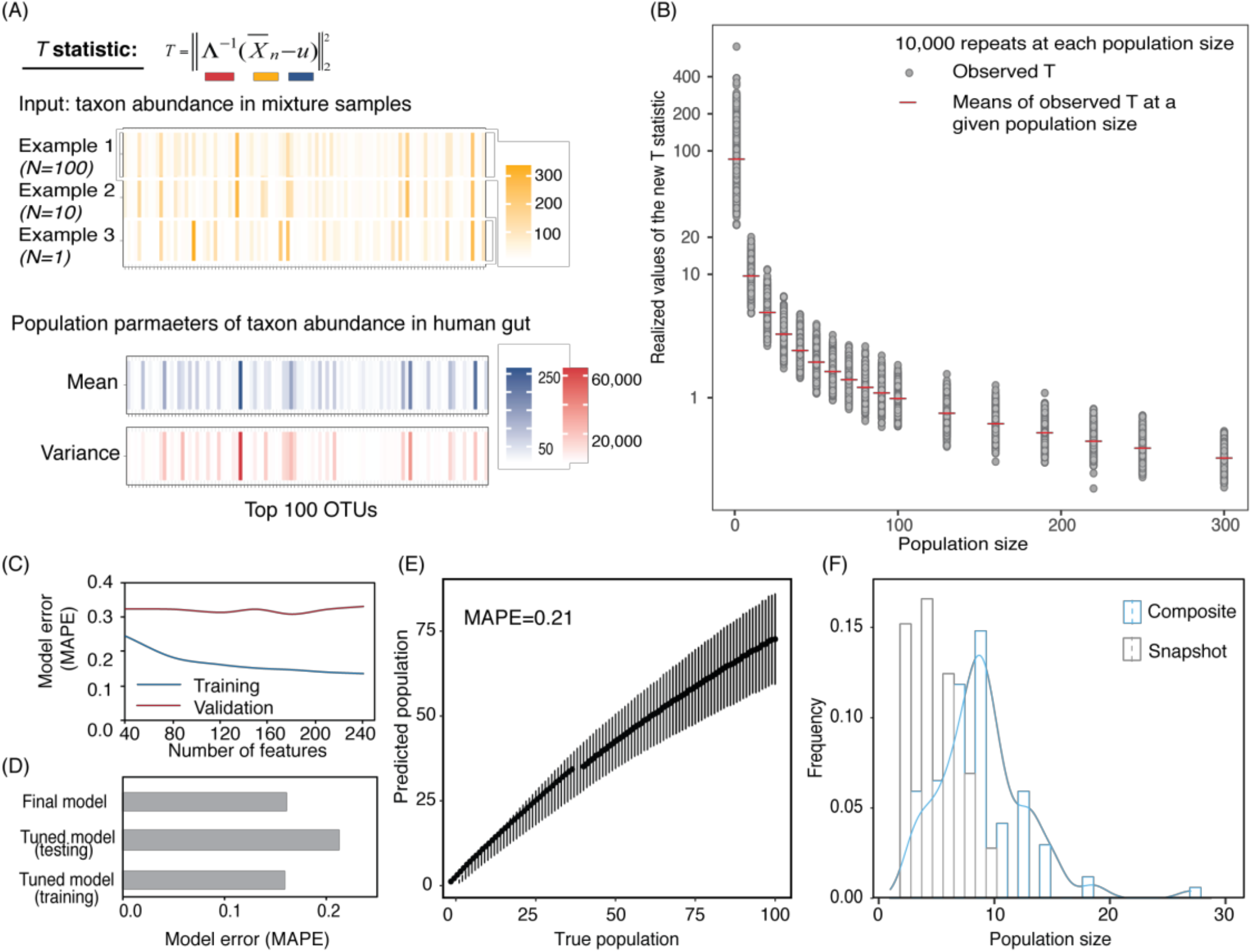
MicrobiomeCensus statistic definition, model training, validation, and application. (A) Example of computing the *T* statistic. (B) Simulation results for *T* with different population sizes. Grey points are simulation results. Red bars are means of 10,000 repeats performed for each population size. (C) Model training and tuning. We built the MicrobiomeCensus model using our *T* statistic and a maximum likelihood procedure. The training set consisted of 10,000 samples for population sizes ranging from 1-300, and 50% of the data were used to train and validate the model. Training and validation errors from different feature subsets are shown. Training errors are shown as red lines, and validation errors are shown as blue lines. (D) Model performance on simulation benchmark. After training and validation, the model utilized the top 120 abundance features. Model performance was tested on synthetic data generated from 550 different subjects not previously seen by the model. The training set consisted of 10,000 samples with population sizes from 1-300, and the testing set consisted of 10,000 repeats at the evaluated population sizes. The training error, testing error, and the error of the final model are shown. (E) Model performance evaluated using a testing set. Black solid dots indicate the means of the predicted values, and error bars indicate the standard deviations of the predicted values. (F) Application of the microbiome population model in sewage. Seventy-six composite samples (blue) were taken from three manholes on the MIT campus, and each sample was taken over 3 hours during the morning peak water usage hours. Twenty-five snapshot samples (grey) were taken using a peristaltic pump for 5 minutes at 1-hour intervals throughout a day.

In developing this new method, we utilize the variance change by population, but without any *priori* assumption about the gut bacterium species taxon abundance distributions and the covariance between species. Our analysis showed that the statistic *T_n_* changed monotonically with increasing population size, indicating the promise of a population estimation model (Figure 3B).

Leveraging our statistic *T_n_*, we constructed an asymptotic maximum likelihood estimator to estimate size of the sample without the information of each individual, that is, we do not observe *X*_1_, …, *X_n_* but only their mixture 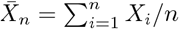. Here, the parameter of interest is the population size n, the test statistic is *T_n_*, and a point estimate is made by maximizing the estimated likelihood of *T_n_* with respect to n. We performed training and validation using 50% of the human microbiome data and held out the rest of the data for testing. Our model achieved a training error as low as 13% (mean absolute percentage error, MAPE) when up to 250 features are included. The model’s training performance increased when more features were included, yet the validation error did not profoundly change with an increasing number of features (Figure 3C). Upon training and validation, we chose the top 120 OTUs and tested the performance of the tuned model on a test set held out during training/validation. The model’s MAPE was 21% (Figure 3D and E, testing errors at each population size evaluated are provided in Table S3), indicating that our model generalized well across different hosts. We then used all data and tuned hyperparameter to acquire a final model. When applying the final model on the same testing data, our model achieved a testing error of 16.2% (Figure 3D).

It is worth noting that in this algorithm, for each size *n*, we only need to estimate the sampling distribution of the statistic *T_n_* once. Hence it is not time-consuming regardless of the true population size. We also note that an RF regression model could not be trained in a reasonable time on the same dataset, even with high-performance computing (Methods). Our model performed remarkably better than a ten-fold cross-validated RF regression model utilizing a reduced dataset, which gave an MAPE of 32%, while the training time for our model was only a fraction of that of the RF regression model (Figure S1).

### MicrobiomeCensus detects human population size differences in sewage samples

With the newly developed population model, we set out to apply our model to sewage samples. Ideally, we would like to apply the model to samples generated from a fully controlled experiment with known human hosts contributing at a given time, yet such an experiment presents logistic challenges. Instead, we applied our model to sewage samples taken using one of two methods, either a snapshot (grab sample) sample taken from the sewage stream over 5 minutes, or an accumulative (composite sample) taken at a constant rate over 3 hours during morning peak human defecation[15] (Figure S2). We hypothesized that the composite samples would represent more people than snapshot samples. Taking grab samples, we sampled at 1-hr intervals at one manhole (n=25); using the accumulative method, we sampled at three campus buildings (classroom, dormitory, and family housing) multiple times over three months (n=76). To remove sequences possibly contributed by the water, we applied a taxonomic filter to retain families associated with the gut microbiome and normalized the species abundance by the retained sequencing reads (Methods, Table S4). We applied our final model to the sewage data set. Our model estimated 1-9 people’s waste was captured by the snapshot samples (mean=3, s.d.=3), and 3-27 people were represented by the composite samples (mean=9, s.d.=7), where the composite samples represented significantly more people (*p* < 0.0001) (Figure 3F). The hypothesis that composite samples represent more people is well supported by our model results.

### Sub-species diversity in sewage samples reflects adding microbiomes from multiple people

Independent from our MicrobiomeCensus model, we found that certain human gut-associated species were frequently detected in sewage samples by using shotgun metagenomics, e.g., *Bacteroides vulgatus, Provotella copri*, and *Eubacterium rectale*. Further, their sub-species diversity, as indicated by nucleotide diversity and the number of polymorphic sites in housekeeping genes, was dramatically higher in sewage samples than in the gut microbiomes of individual human subjects (Figure 4A-F and Supplementary Results).

**Figure 4.**
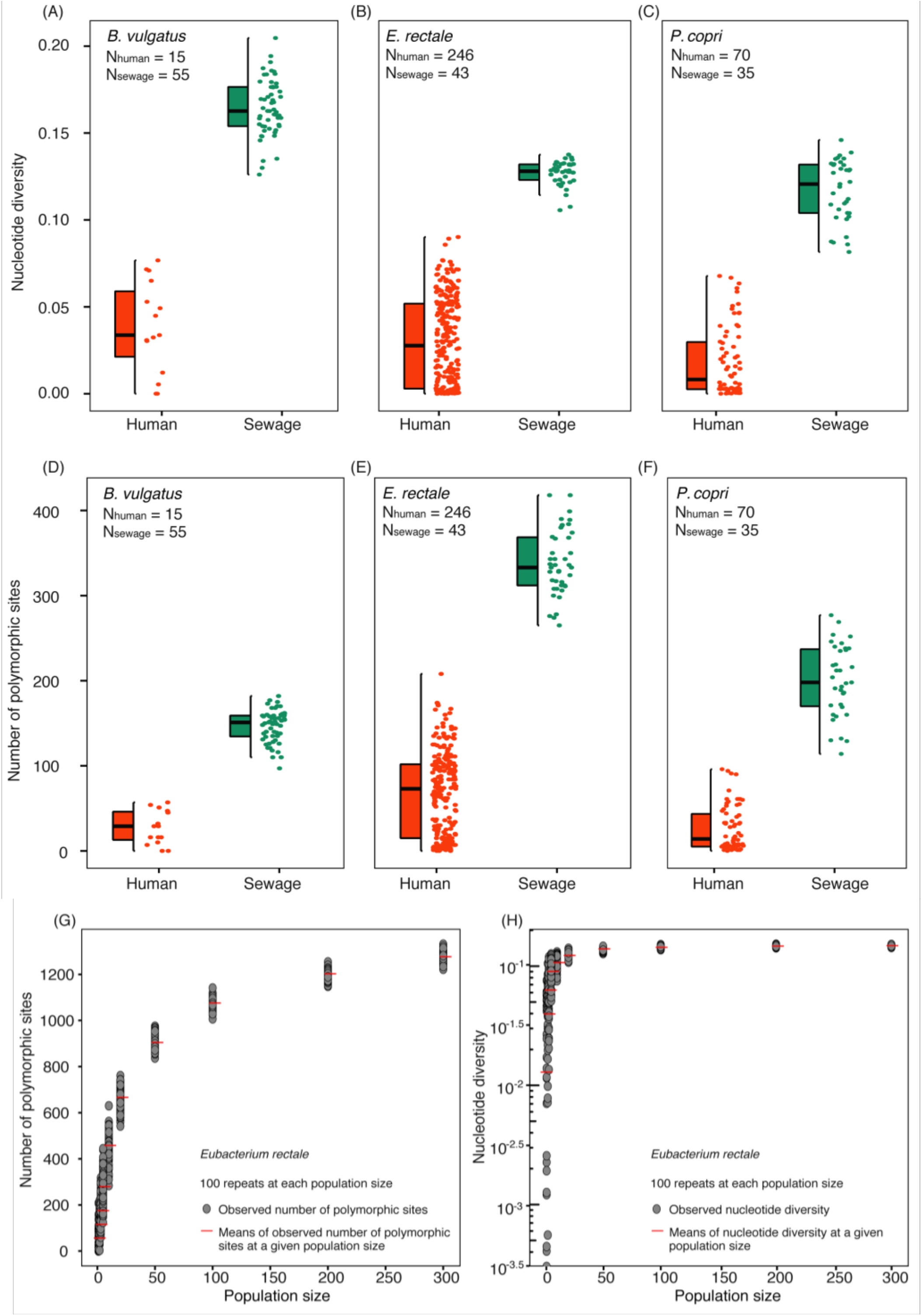
Sub-species diversity in gut-associated bacterial species as a potential marker for human population size. (A-F) Comparison of sub-species diversity of gut-associated bacteria in human gut microbiome samples (LifelinesDeep) and MIT sewage samples. Nucleotide diversity and numbers of polymorphic sites were computed from ten phylogenetic marker genes. (G) and (H) Simulation results showing intra-species diversity in response to increasing population size, as represented by the number of polymorphic sites (G) and nucleotide diversity (H).

To examine the effect of increasing population size on sub-species genetic variation in representative gut-associated microbial species, we simulated aggregate human gut samples using a sample without replacement procedure and computed the nucleotide diversity and numbers of polymorphic sites for the aggregate samples at different population sizes. This resulted in SNV profiles from 64 species. Our simulation showed increases in both nucleotide diversity and the number of polymorphic sites as more human gut samples were aggregated (Figure 3 G and H). For instance, the nucleotide diversity and number of polymorphic sites in *Eubacterium rectale* increased from 0.029 (s.d. 0.026) to 0.149 (s.d. 0.002) and 64 (s.d. 54.33) to 1274 (s.d. 18.41), respectively, when the population size increased from 1 to 300. Further, the number of polymorphic sites strongly correlated with the population size (Pearson correlation coefficient > 0.8) in 49 of the 64 species (Table S7), suggesting the potential that the SNV profiles of a wide range of gut species could be developed into feature space for population size estimation. Our simulation further shows that the number of polymorphic sites increased with population size more slowly than nucleotide diversity, indicating its potential to reflect more subtle changes in population size (Figure 4G and H). Despite the need for further model developments, the analysis here shows the potential of the sub-species diversity of gut anaerobes as a feature space to be developed into a population size estimation model, independent from the taxon abundance-based model described here.

## Discussion

The MicrobiomeCensus method we present here can, in theory, estimate the population size contributing to a sewage sample from the taxon abundance of multiple human gut microbiome taxa, using our T statistic and associated maximum likelihood estimation and application procedures. While the model is trained to perform accurate population estimation on a neighborhood scale, we expect the population range it can estimate to expand with increasing training gut microbiome data availability. We propose the MicrobiomeCensus model as a tool to drive further developments in quantitative sewage-based epidemiology. We have provided mathematical proof of the validity of our approach.

MicrobiomeCensus showed excellent performance in our simulation benchmark. In particular, the study subjects that we utilized in the training and testing sets are random samples out of 1,100 men and women across a wide range of age without any stratification, hence the model’s testing performance indicates its generalizability. Our study is founded on the observations that healthy gut microbiomes are resilient, with inter-individual variability outweighing variability within individuals over time [12, 20, 26]. There are caveats to our approach; potentially, diets and regional effects on human microbiome composition could introduce noises to the prediction [17, 14]. In applications to sewage, future studies on water matrix effects should be performed to understand and further account for noises from the sewage collection network. It should be mentioned that while our model is trained on microbiome data, it is not limited to microbiome features because we did not impose any assumptions on the distribution of features. Other features, e.g., crAssphage titers, may also be incorporated once individual level data at a large population size become available to allow model validation.

Utilizing sewage to understand population-level dynamics of human symbionts presents a new scenario of sampling meta-communities. The gut microbiomes of humans can be viewed as local communities, and gut microbiomes of people living in a neighborhood could be viewed as a kind of regional meta-communities, because these communities are linked by dispersal that can take place among people connected by social networks and through a shared built environment. The meta-community framework is considered to provide useful new conceptual tools to understand the largely unexplained inter-personal variability in gut microbiomes, with expansions of the theory to consider biotic interactions suggested by Miller, Svanbäck, and Bohannan [27]. In considering a sample of meta-communities, Leibold and Chase asked provocatively “what is a community?” and observed that the definition of a community is usually “user-defined and could be context-dependent” –”one community ecologist might explore the patterns of coexistence and species interactions among species within a delimited area, the other might ask the same question but define a community that encompasses more area and thus types of species, as well as different degrees of movements and heterogeneity patterns” [19]. The ambiguity between samples of meta-communities and local communities is particularly challenging for samples of microbial communities, because dispersal boundaries are difficult to delineate. Despite the conceptual importance, empirical methods that explicitly test whether a microbiome sample is a sample of a meta-community or a local community have not been available. MicrobiomeCensus directly distinguishes samples of meta-communities and local communities by enumerating the number of hosts contributing to a microbiome. While MicrobiomeCensus is trained on gut microbiome data, the procedure may have wide applications in other microbial ecosystems.

In response to the COVID-19 pandemic now affecting the human population globally, sewage-based virus monitoring is underway[3]. Our analysis calls for attention to the denominator used in normalizing the biomarker measurements. While in practice, loading-based population proxies such as the copy numbers of pepper mild mottle viruses are used to normalize data generated from sewage, such proxies would likely have high error at the neighborhood level because of their variability in human fecal viromes (10^6^-10 virions per gram of dry weight in fecal matters)[38], while they likely have reasonable performance when the population size is sufficiently large and the means of biomarker loadings converge under the Central Limit Theorem. Thus, the relationships between sewage measurements and true viral prevalence in small populations are hard to establish despite the need for sentinel population studies. Our model has immediate application in detecting false negatives, because it alerts us to the possibility that an absence of biomarkers might be caused by a sewage sample that under-represents a population. With further developments incorporating local training data, the model can potentially generate a denominator that can help turn biomarker measurements into estimates of prevalence and enable the application of epidemiology models at finer spatio-temporal resolutions.

## Methods

### Rationales

If the distribution of *X_i_* is known, then the distribution *T_n_* is known and one can easily use maximum likelihood estimator (MLE) to estimate the human population size. Here the human population size is essentially the sample size n from the wastewater sample. However for generality, we do not want to impose any specific distribution assumptions on taxon abundance distributions, thus, we need to rely on asymptotic results to estimate the distribution of our statistic. Unlike the univariate case where the asymptotic distribution of the statistic *T_n_* can be simply derived by central limit theorem, we are dealing with a much more challenging situation due to high dimensionality.

In the following, we shall firstly introduce some notations and assumptions that will be needed for the theorems. Then, in Theorem 1, we derive the Gaussian approximation for the test statistic *T_n_*, which implies that we can use simulated Gaussian vectors to approximate the distribution of our statistic. To apply this approximation, one needs to further estimate the covariance matrix which is highly non-trivial due to high dimensionality. To get around this difficulty, we further propose a sub-sampling approach. Theorem 2 provides the theoretical foundation for this sub-sampling scheme.

### Notation

Recall that vectors *X*_1_, *X*_2_, …, *X_n_* are microbiome profiles from individuals 1, 2, …, *n*. Assume that they are independent and identically distributed (i.i.d) random vectors in 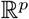 with mean 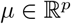 and variance 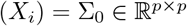. Denote 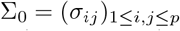 and 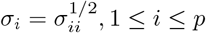. Let the diagonal matrix Λ_0_ = diag(*σ_i_*, 1 ≤ *i* ≤ *p*).

To construct the Gaussian approximation, we shall firstly work with cases when both Λ_0_ and *μ* are given, that is, consider the statistic

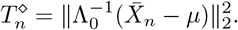

We will later extend all the results to cases when those parameters are unknown. For notation’s simplicity, consider the normalized version of *X_i_*:

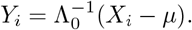

Then 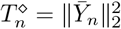, where 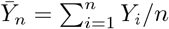, and the covariance matrix ∑_*Y*_ of *Y_i_* is the correlation matrix of *X_i_*, with expression 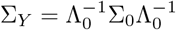.

We need the following condition on *Y_i_* for the main theorem.

#### Assumption 1

*Let s* > 2. *Assume*

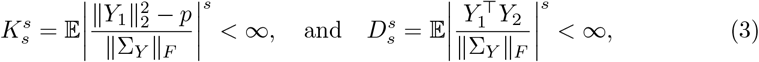

*where* ||.||_*F*_ *is the matrix Frobenius norm*.

Remark 1 *Above conditions naturally hold if Y*_1*i*_, 1 ≤ *i* ≤ *p*, *are independent and* 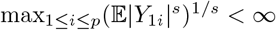. *Actually under this setting, ∑_Y_* = *I_p_ and thus* ||∑_Y_||_*F*_ = *p*^1/2^. *By Berkholder[6]’s inequality*,

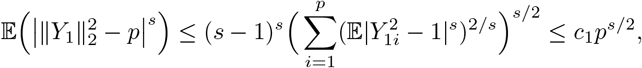

*where c*_1_ > 0 *is independent of p. This justifies K_s_ part in condition* (3). *Again by Berkholder[6]’s inequality*,

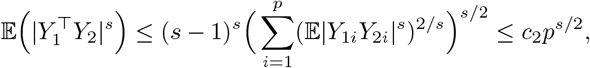

*where c*_2_ > 0 *is independent of p. And thus D_s_ part in condition* (3) *holds*.

Now we are ready to introduce the first asymptotic result. The following theorem essentially states that under certain regularity conditions, the distribution of our test statistic can be approximated by the distribution of some function of a Gaussian vector. T

#### Theorem 1 (Theorem 2.2 in Xu et al.[36])

*Assume Assumption 1 holds with some s > 2, also assume*

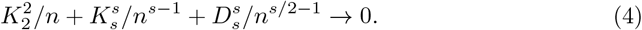

*Then for Z* ~ *N* (0, ∑*_Y_*), *we have*

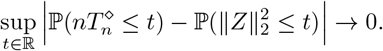

It is worth noticing that under settings in Remark 1, condition (4) holds. Based on the above theorem, if we can estimate the covariance matrix ∑_*Y*_, then 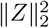 can be generated and used for approximation of our statistic. However the estimation of ∑_*Y*_ is difficult due to high dimensionality unless some additional conditions are imposed on the covariance matrix. To get around this difficulty, we shall consider a bootstrap approach. The main idea is that for each *n*, we randomly generate *n* samples from the reference data *X_i_*, 1 ≤ *i* ≤ *n*_0_, and construct some generated statistic. Using the empirical distribution of the generated statistic to approximate the distribution of our statistic. In the following, we shall provide the theoretical justification for this procedure.

For some integer *J* > 0, let *A*_1_, *A*_2_, …, *A_J_* be i.i.d uniformly sampled from the class 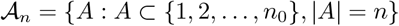. Assume the sampling process are independent from our data (*X_i_*)_*i*_. Then for each 1 ≤ *k* ≤ *J*, the set {*X_i_, i* ∈ *A_k_*} is of size *n* and can be used to construct one test statistic 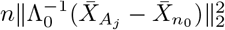, where 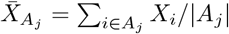. After repeating this procedure *J* times, we would have *J* realizations of our test statistic which can be used to construct the empirical distribution 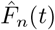:

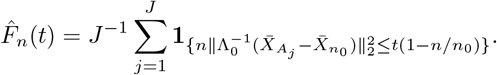

We shall later show that this empirical distribution can be adopted for approximating the distribution of the target statistic 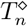. Following result can be derived based on Theorem 3.5 in Xu et al.[36] and Theorem 1.

#### Theorem 2

*Assume conditions in Theorem 1 hold, and moreover assume that n* → ∞, *n* = *o*(*n*_0_) *and* (4) *holds. Then for J* → ∞, *we have*

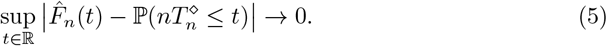

Theorem 2 implies that we can use the empirical distribution generated from our sub-samples to estimate the distribution of our target. As mentioned previously, so far, we are assuming that we know the value of Λ_0_ which is not realistic in applications. Therefore we need to further estimate Λ_0_, that is, we need to estimate the standard deviation for each coordinate. This can be easily accomplished by considering the estimator

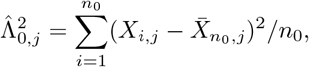

where *X_i,j_* represents the *j*th coordinate of *X_i_*, and Λ_0, *j*_ is the jth diagonal entity of Λ_0_ and *n*_0_ is the size of the reference data. Also one can use the average of the reference data to replace *μ*. If *X_i,j_* has heavy tail, we can also consider a robust m-estimator for ∑_0_ and *μ*, see for example, Catoni[8].

### Bootstrap procedure

Below we describe the bootstrap procedure we use to approximate the distribution of *T_n_* for different census counts. Recall that *X*_1_, …, *X_n_* represent arrays of taxon relative abundances in the gut microbiome of human subject 1, …, *n*, and *T_n_* is defined in (Eq. 2).

Step 1. Estimate the population mean 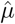, and the diagonal matrix 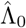, from a reference sample human gut microbiome data.
Step 2. For each census count *n*, generate 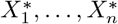 from the reference data to compute 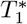.
Step 3. Repeat Step 2 *B* times (herein 10,000 times) to get 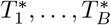.
Step 4. Obtain the density function 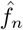 of *T_n_* based on 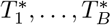 using a Gaussian kernel.
Step 5. Repeat Steps 2-4 for all the census counts *n* =1, 2, …, *N* considered, herein *N* = 300. It should be noted that per Theorem 2 we require bootstrap sample size *n* much smaller than total reference sample size *n*_0_, thus up to 300-person samples were simulated here because the gut microbiome reference dataset we utilized consisted of a total of 1,100 people. The range can be expanded if a larger dataset is available.

### Maximum Likelihood Estimation

We use a maximum likelihood estimation (MLE) procedure to achieve point estimates of the population size from a new mixture sample, *W*. The MLE procedure firstly computes *T*_0_ by replacing the sample average by *W*, that is

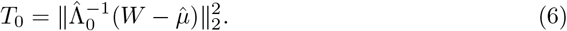

And then computes the likelihoods that *T*_0_ was drawn from population sizes from 1 to *N*, respectively, using the estimated distributions generated from the bootstrap procedures described above. Next, the population size 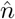 that yields the highest likelihood is chosen.

### Confidence interval

Due to Theorem 1, the asymptotic distribution of *nT_n_* is the same as 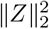 and is therefore independent of the parameter *n*. Hence for any confidence level 1 – *α*, we can firstly estimate the 1 – *α* quantile of *nT_n_* based on the simulated data described above, denoted as 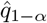. Then for any new mixture *W* and the corresponding *T*_0_ as in (6). We our 1 – *α* confidence interval is 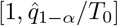.

### Model training, validation, and testing

We synthesized a mixed data set from a gut microbiome dataset of 1,135 healthy human hosts from the Lifelines Deep study[39], which was the largest single-center study of population-level human microbiome variations from a single sequencing center at the time of this study. The data set consisted of 661 women and 474 men. We considered OTUs defined by 99% similarity of partial ribosomal RNA gene sequences (Methods of OTU clustering are described in detail in Supplementary Methods). After quality filtering, we retained 1,100 samples that had more than 4,000 sequencing reads/sample. We split the entire dataset approximately in half, using 550 subjects to generate the training/validation set and the other 550 subjects to generate the test set. We then used the aforementioned ideal sewage mixture approach to generate synthetic populations of up to 300 individuals, which is the relevant range for population estimation in upstream sewage. The training error was computed using the entire training data set. Five repeated holdout validations using a 50-50 split in the training set were performed to tune the hyperparameter for feature selection. The training and cross-validation errors were evaluated at integers from 1 to 100, using the error definition:

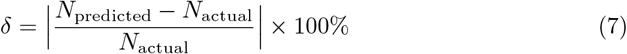

and the model’s performance across all the population sizes was characterized by the mean absolute percentage error (MAPE):

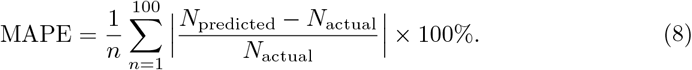

We chose to use MAPE because for a problem of population estimation, the the error relative to the true value is important to consider for performance evaluation. After training and validation, the hyperparameter (in this case, the top *k* abundant OTUs) that yielded the best performance in the validation step was used in the model. The tuned model was then tested on the test set. Our synthetic sewage microbiome approach captured the actual microbiome variation among individual hosts and demonstrated the model’s generalizability.

### Human gut microbiome 16S rRNA amplicon data source

The single-person and multi-person microbiome data were drawn from a gut microbiome dataset of 1,135 healthy human hosts from the Lifelines Deep study [39], which was the largest study of population-level human microbiome variations from a single sequencing center at the time of this study. The data set consisted of 661 women and 474 men. We considered operational taxonomic units defined by 99% similarity of partial ribosomal RNA gene sequences. After quality filtering, we retained 1,100 samples that had more than 4,000 sequencing reads/sample. The rarefaction depth was chosen to balance sample size and sequencing depth.

### 16S rRNA gene amplicon sequencing data analysis

Operational taxonomic units defined at 99% sequencing similarity were generated from the combined dataset by first denoising the samples with DADA2[7], and then clustering the outputted exact sequence variants with the q2-vsearch plugin of QIIME2 [33]. Taxonomic assignments were performed using a multinomial naïve Bayes classifier against SILVA 132[32, 4]. All 16S rRNA gene amplicon analyses were performed in the QIIME2 platform (QIIME 2019.10) [5].

### Species Abundance Distribution

We examined the relationships between the performances of several widely used SAD models and the number of contributors (population size) to a multi-person microbiome. Multi-person microbiomes were generated by sampling N individuals from the quality-filtered gut microbiome 16S rRNA dataset and summing the abundances of the same taxa. At each population size, 10,000 repeats were performed. The repeats were chosen according to the constraints of computational efficiency. The SADs evaluated included the Lognormal, Poisson Lognormal, Broken-stick, Log series and the Zipf model, which were shown to have varied successes in predicting microbial SADs [34]. We examined the fit using a rank-by-rank approach as previously described by Shoemaker et al. [34]. First, maximum-likelihood coefficients for each of the SADs described above were estimated using the R package sads[30]. Next, SADs were predicted using each model, and tabulated as RADs. Then, we used a least-squares regression to assess the relationship between the performance of the predicted SADs against the observations and recorded the coefficient of determination (R-squared). Last, R-squared values from model fits of each SAD model were summarized as the means, and the models that resulted in the highest R-squared values for each simulated community were recorded.

### Field data

We conducted a field sampling campaign, collecting sewage samples daily at manholes near three buildings (two dormitory buildings and one office building) on the campus of Massachusetts Institute of Technology. Seventy-six sewage samples were collected through a continuous peristaltic pump sampler operated at the morning peak (7-10 a.m. near the dormitory buildings and 8-11 a.m. near the office building) at 4 mL/min for 3 hours. Wastewater was filtered through sterile 0.22-μm mixed cellulose filters to collect microbial biomass. Environmental DNA was extracted with a Qiagen PowerSoil DNA extraction kit according to the manufacturer’s protocol. The DNA was amplified for the V4 region of the 16S rRNA gene and sequenced in a Miseq paired-end format at the MIT BioMicro Center, according to a previously published protocol[31]. Included as a comparison are a set of snapshot sewage samples taken using a peristaltic pump sampler at 100 mL/min for 5 minutes over a day (10 a.m. on Wednesday April 8, 2015, to 9 a.m. on Thursday April 9, 2015). The sampling methods for snapshot samples are described in detail by Matus et al.[24]

### Application to sewage data

1he 16S rRNA gene amplicon sequencing data from the field sewage samples were trimmed to the same region, 16S V4 (534-786) with the LifeLines Deep data using Cutadapt 1.12 [23]. Forward reads were trimmed to 175bps, and reverse reads were first trimmed to 175bps and then further trimmed to 155bps during quality screening. We created a taxonomic filter based on the composition of the gut microbiome data set, which consisted of the abundant family-level taxa that accounted for 99% of the sequencing reads in the human gut microbiome data set, and excluded those that might have an ecological niche in tap water (*Enterobacteriaceae* and *Burkholderiaceae*). This exclusion resulted in 25 bacterial families and one archaeal family in our taxonomic filter, including *Lachnospiraceae*, *Ruminococcaceae*, *Bifidobacteriaceae, Erysipelotrichaceae, Bacteroidaceae*, and others (Table S4). We applied our taxonomic filter to the sewage sequencing data, which retained 73.9% of the sequencing reads. This retention rate is consistent with our previous report of the human microbiome fraction in residential sewage samples[24]. We then normalized the relative abundance of taxa against the remaining sequencing reads in each sample. Welch’s two-sample t-tests were performed to retain the OTUs whose means did not differ significantly from the human microbiome data set (p > 0.05).

### Deployment of generic machine learning models

Logistic regression, support vector machine, and random forest classifiers were employed to perform the classification task for population sizes of 1, 10, and 100. Model training, cross-validation, and testing were performed using the R Caret platform with the default setting[18]. For the support vector machine, the radial basis function kernel was employed. Ten-fold cross-validation and five repeats were performed for all the models considered. Model performance was evaluated using accuracy, sensitivity, and specificity. Based on the classifier performance, the RF regression model was used for comparison with our new model’s performance. Initially, we trained the model using the same training data set used in training our maximum likelihood model, however, the computation was infeasible, even with a 36-thread, 3TB-memory computing cluster. We then introduced gaps in the population size range, using populations from the vector (1, 5, 10, 20, 30, 40, 50, 60, 70, 80, 90, 100, 110, 120, 150, 180, 240, 300)^⊤^ while maintaining the same sample size at each population size (10,000 samples). The training was performed in R Caret, using 10-fold cross-validation. Ten variables were randomly sampled as candidates at each split, mtry=10. The performance was evaluated using the same testing set that was used to evaluate the maximum likelihood model.

## Acknowledgements

We thank Cho C. Yiu, David E Hingston, and Joseph S. Monteiro from the MIT Facilities department for assistance with sewage sampling. We thank Noriko Endo, Sean Gibbons, Tami Lieberman, Xiaofang Jiang and Shijie Zhao for valuable discussions. We thank Mariana Matus and Newsha Ghaeli for acquisition and access of 24-hr sewage time series data. The sampling and sequencing experiments were performed with funding from the Kuwait Foundation for Advancement of Sciences. Fangqiong Ling’s work at MIT was partially supported by an Alfred P. Sloan Foundation Microbiology of the Built Environment Postdoctoral Fellowship. The computational analyses were performed with resources provided by the WUSTL McKelvey School of Engineering. Lin Zhang was supported by National Science Foundation Grant 1322-58679 awarded to Fangqiong Ling.

## Author contributions statement

F.L. and E.J.A. designed the study; F.L. performed simulation; L.C. performed mathematical proofs; F.L., X.Y. and L.Z. performed sequence analysis; F.L., S.I., C.Dai., and S.P. performed sewage experiments; C. Duvallet, K. F-M, F.D. C.R. and E.J.A. coordinated the acquisition of sequencing data; L.Z., L.C., E.J.A and F.L. wrote the manuscript.

## Code and data availability

Source code will be made avaiable through https://github.com/linglab-washu/population-model upon publication. Sewage metagenomic data will be made available at National Center for Biotechnology Information Short Read Archive at BioProject PRJNA683921 upon the time of publication.

